# Efficient Stochastic Trace Generation for Transcription^⋆^

**DOI:** 10.64898/2026.05.05.722871

**Authors:** Arman Ferdowsi, Matthias Függer, Thomas Nowak

**Author notes:** This is the bioRxiv version of the paper accepted at CMSB’26, to appear in Lecture Notes in Bioinformatics.

## Abstract

Bursty transcription in single cells typically produces over-dispersed, skewed, and sometimes heavy-tailed expression distributions that are explained by two-state Markov models of the promoters. While the gold standard for simulation is exact stochastic sampling with Gillespie’s algorithm, obtaining thousands of timed traces is computationally costly. Surrogate models based on stochastic differential equations (SDEs) are widely used to speed up this simulation process. An example is the Chemical Langevin Equation based on Gaussian noise, which, however, does not capture heavy-tailed noise.

In this work, we present a unified SDE framework that combines deterministic drift, Gaussian fluctuations, and additive sporadic jumps of arbitrary distributions, and provide an open-source Python implementation, `bcrnnoise`. The framework subsumes standard surrogate models and allows for vectorized generation of batches of transcription traces. We assess computational speed and accuracy of common surrogate models along with new models, showing that high accuracy can be obtained while reducing computational cost up to two orders of magnitude.

## 1 Introduction

Many biochemical processes show pronounced stochastic behavior at different levels of abstraction. One such process happening in every cell is transcription, the process of producing mRNA from a DNA sequence. Single-cell measurements have repeatedly shown that gene transcription often occurs in episodic bursts, where mRNA is transcribed, followed by pauses of no activity [41]. Direct imaging and sequencing-based assays have quantified burst frequency and size at transcript resolution, revealing behavior that aligns well with a promoter probabilistically toggling between two states [35,36,41]: an active on state where transcription occurs with a certain rate and an off state with no transcription. This so-called two-state telegraph transcription model is widely used [27,33,41,34,26] to capture transcription bursts, an effect that is not explained by the simpler, also commonly used, one-state (constitutive) model [39,32], where transcription occurs at a constant rate.

Such a biochemical system can be readily described within the framework of Chemical Reaction Networks (CRNs) [13,38], where biochemical species interact with each other via reactions. The resulting dynamics *p*(*x, t*) of the probability of the species counts being equal to vector *x* at time *t* are described by the Chemical Master Equation (CME). The CME is rarely directly used for the generation of traces. The gold standard to numerically solve such models is exact stochastic sampling via Gillespie’s algorithm [13]. Computationally, this is expensive even for systems with only a few species like the two-state transcription model. In particular, high event rates, e.g., due to high plasmid copy number and thus many genes being transcribed, lead to slow progress in simulated time, since the state vector is updated at each event. Furthermore, thousands of simulations have to be run to obtain detailed noise profiles of the stochastic process. Indeed, detailed noise characteristics have been used for model discrimination in single-molecule assays, by comparing predicted and observed noise profiles to reject incompatible mechanistic hypotheses and guide experimental design [18].

To address this limitation, computationally faster algorithms are widely used and have been implemented in software packages such as COPASI [20], BasiCO [3], and StochKit [37] the simulation of arbitrary CRNs. Prominent algorithms are discrete-noise approximations such as Tau-leaping [15] where several events are grouped together before updating the system’s state and continuous-state surrogates based on Stochastic Differential Equations (SDEs). The Chemical Langevin Equation (CLE) is an example of the latter, replacing reaction-counting fluctuations by Gaussian noise, yielding tractable SDEs for means, variances, spectra, and control-theoretic analyses [12,14,38,24]. While fast, surrogates like the CLE fail to capture the heavy tails and overdispersion characteristic of processes like burst-driven transcription [11,29,8,35,31] and are thus less informative for model discrimination.

Abstract models that account for non-Gaussian behavior have been proposed in the context of transcription. A promising approach is replacing parts of the transcription process with piecewise-deterministic processes [29], which showed improved accuracy on steady-states compared to CLE models. Also, models that study the case of increasing volume *V→*∞with Lévy behavior have been proposed as abstract models [23]. However, fast surrogate models that remain accurate over the complete time trajectory for small cellular volumes and that account for transient effects after activating transcription remain challenging.

Related to this line of work is also analytical progress that has been made for transcription models. For the two-state telegraph model, analytical results are available in generating-function form and yield closed-form moment expressions [34,21], providing an important reference for marginal distributions. However, the work does not explicitly address multi-copy settings as they occur, e.g., in bacteria where a plasmid with a certain copy number carries identical DNA sequences from which transcription occurs.

In particular, activating promoters leads to a transient behavior from all promoters being in the off state towards random switching of promoters. Further, these analytic results do not directly enable fast generation of complete traces since they a priori lack fast transition kernels, and correlations between different times, needed for large-scale trace generation and noise-profile comparisons.

Of mention is also research that builds on analytical progress of such models, aiming towards more detailed transcription models beyond the two-state model. Examples are work on models that include other cellular processes for which approximate analytic solutions have been obtained [5], an approximate analytic model for gene expression with stochastic bursts [28] that also includes the translation process from mRNA to protein and a comparison to commonly used diffusion-type models, and an analytic model that includes splicing [17].

### Contributions

We propose a unified Lévy–Itô jump-diffusion SDE framework that places common CRN approximate models like Tau-leaping and CLE along compound-Poisson burst surrogates within a single drift–noise specification. Compound-Poisson and, more generally, Lévy jump processes can replace or complement Gaussian diffusion noise terms in SDEs, enabling heavy tails [1,7]. We implemented the SDE framework, making it available as an open-source Python library `bcrnnoise` that provides a common simulation API for physical-unit-aware specification of deterministic, diffusion, and jump processes. The library enables vectorized simulation of large numbers of traces within batches in discrete update steps of some specified time step *Δt >* 0, in contrast to sequential simulation of such traces one by one. While we use natively compiled libraries and just-in-time-compilation techniques to speed up the Python code, the focus of this work is on the systematic comparison of noise models for transcription rather than highly optimized CRN simulation.

For comparison with gold standard simulations, the library also implements Gillespie’s algorithm.

We compare common and newly proposed surrogate models for the one-state (constitutive) and the two-state (telegraph promoter) transcription models within our framework for computational time and accuracy of their noise profile. For parameterization of the surrogate models we derive analytic formulas from stationary moments (Theorem 1). Our simulations confirm that, for low plasmid copy numbers, CLE approximations of the two-state model show low accuracy. Further, models based on Geometric noise perform relatively well; in particular, if in variants accounting for transient behavior. Finally, a Binomial-Poisson noise model is shown to outperform the widely-used Tau-leaping algorithm for large update-steps *Δt >* 0 in terms of accuracy, rendering it a promising candidate for vectorized generation of high numbers of traces within short time at high accuracy: our simulations show up to two orders of magnitude improvement in runtime with high accuracy.

## 2 Canonical Stochastic Transcription CRNs

We consider the two most-used CRN models of transcription, differing in whether the promoter is constitutively active or switches between discrete activity states, resulting in transcription bursts. We further assume that the gene is present *n*_p_ times within a fixed volume *V* of a cell, e.g., due to the copy number of the gene-bearing plasmid being *n*_p_.

### Constitutive one-state model

Each plasmid constitutively produces mRNA at rate *α* (min^−1^ fL^−1^), giving a total transcription rate of *αn*_p_. Transcripts are degraded or diluted at first-order rate *δ* (min^−1^). The resulting CRN is given as

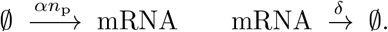

We assume that initially there is no mRNA in the system.

The exact stationary distribution of the mRNA copy number under this CRN is Poisson with mean *αn*_p_*V/δ* [39,32] (Supplementary Material, Theorem 2).

### Telegraph two-state model

Each plasmid independently switches between a transcriptionally inactive state (off) and an active state (on) at rates *k*_on_ and *k*_off_ (min^−1^), respectively. Only on plasmids produce mRNA, at rate *α* (min^−1^) per active plasmid, and transcripts are degraded as above. Denoting concentrations of on and off plasmids by *N*_on_ and *N*_off_, with conservation law *N*_on_ + *N*_off_ = *n*_p_*/V*, the CRN is

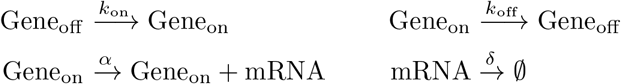

We assume that initially there is no mRNA in the system and that all genes are in the off state. The latter models the scenario where transcription of a gene is activated that is located on multiple copies of a plasmid.

The telegraph model is seen to capture promoter bursting: if *k*_off_ *α* the state is short-lived and mRNA is transcribed in bursts. The exact stationary distribution of mRNA is a Poisson mixture whose mean and variance admit closed-form expressions [34,35] (Supplementary Material, Theorem 3).

### Exact Simulation

Both CRNs are simulated exactly via Gillespie’s stochastic simulation algorithm (SSA) in `bcrnnoise,` which samples each reaction event in continuous time. The one-state model (ℳ_CME_) tracks a single variable (mRNA count) and fires two reactions (birth and death). The two-state model (ℳ _CME,2_) tracks three variables (mRNA count and the numbers of on/off promoters) and fires four reactions. In both cases the per-step time (i.e., the inter-event time) is variable and the simulation cost grows with the total reaction rate, which scales with *n*_*p*_. Benchmarks confirm the expected linear scaling in *n*_*p*_ (Fig. 1a&b).

**Fig. 1:**
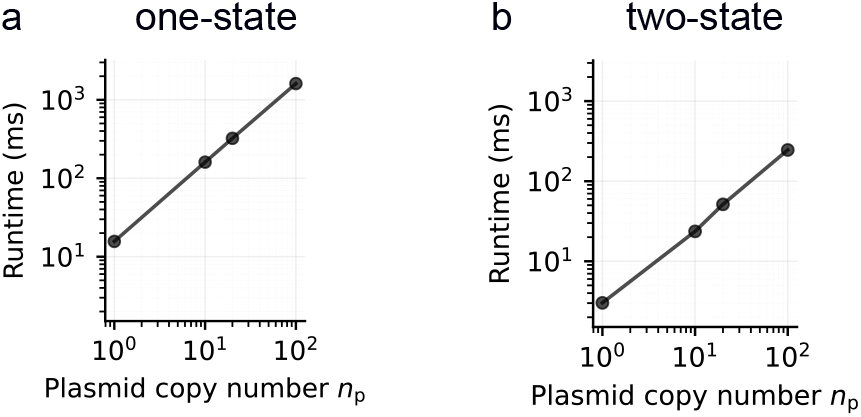
Gillespie SSA runtime scales linearly with plasmid copy number. Simulation time per trajectory (simulated time 120 min) with Gillespie SSA in `bcrnnoise` for the constitutive one-state model (**a**) and the telegraph two-state model (**b**). Means over 20 runs for *n*_*p*_ from 1 to 100 are shown. Parameters follow experimentally reported parameters [4] (one-state) and [16] (two-state); see Methods for details.

Besides the linear scaling of cost per generated trajectory, simulation of several trajectories is a priori not vectorized and may thus be parallelized only over the number of computing cores.

## 3 Unified Jump-Diffusion Framework

We next introduce a unified SDE framework to express surrogate noise models for transcription. Formalizing them within a unified framework allows us to directly compare models. Further, advantages are ease of switching and prototyping noise models within the corresponding Python implementation `bcrnnoise`.

### 3.1 SDE Model

Consider a CRN with *n*≥ 1 biochemical species and let *X*_*t*_ be the stochastic *n*-dimensional vector of the species’ concentrations at time *t*≥ 0. Combining deterministic kinetics, Gaussian fluctuations, and sporadic jumps into a Lévy–Itô equation, one obtains

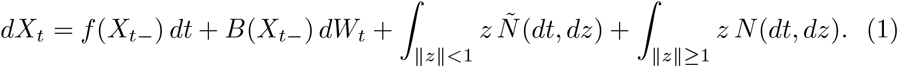

Here *f* is the deterministic drift, *B dW*_*t*_ the diffusive Gaussian component, and the jump terms encode arrival times and jump sizes through the Lévy measure *Π*. One can show that such a combined process is mathematically well-behaved: under mild conditions, it has a unique solution and remains positive in presence of certain boundary conditions for noise (Supplementary Material, Theorem 4). Further, analogous to the CME, analytic equations for the density *p*(*x, t*) over species concentration vectors *x* and time *t* can be stated (Supplementary Material, Theorem 5). In general, these are intractable to solve analytically, however; much like for the CME.

The Tau-leaping and CLE approximations of the CME are recovered as special cases of Eq. (1). *Tau-leaping* sets *B*≡ 0 and takes *Π* to be the reaction-channel jump measure: for each reaction *j* with propensity *a*_*j*_ (*x*) and stoichiometric change *v*_*j*_, one Poisson random measure *N*_*j*_ contributes jumps of size *v*_*j*_ */V* at rate *a*_*j*_ (*x*)*V*, yielding a pure-jump process without Gaussian noise and deterministic components (a pure-jump process without Gaussian noise). *CLE* instead sets the jump dynamics to *Π*≡ 0 and replaces each Poisson channel by a Gaussian with the same infinitesimal mean and variance, giving the state-dependent diffusion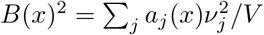.

#### Linear one-dimensional transcription model

While not all surrogate models discussed in this work are one-dimensional, a lower dimensionality of the system’s state is of interest towards the goal of a computationally efficient instance of the unified SDE model. We thus consider the ansatz of a linear (Ornstein– Uhlenbeck-type) one-dimensional model for which we will derive moment expressions that will enable parameterization of this simple model via moment-matching to the full model.

The Ornstein–Uhlenbeck-type model is obtained from (1) with a single dimension, where *x* is the concentration of mRNA, and letting *f* (*x*) = *λ*_0_ − *δx* with transcription rate *λ*_0_ ≥0 and degradation/dilution rate *δ >* 0. Further, the process *W*_*t*_ is a standard Brownian motion, *N* (*dt, dz*) is a Poisson random measure with Lévy measure *Π*, and *Ñ* (*dt, dz*) denotes the compensated measure on {0 *< z <* 1 }. In the compound-Poisson case, *Π*(*dz*) = *κv*(*dz*), so bursts arrive at rate *κ* with jump-size law *Y ∼v*. In fact, letting 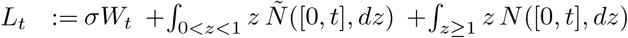 the stationary solution can be written explicitly as 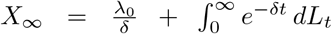, and moments of *X*_∞_ exist precisely up to the corresponding moments of the Lévy input *L*_1_. A derivation of this representation and the corresponding stationary transform formulas is given in Supplementary Material, Section A.4.

Here, we focus on the compound-Poisson (finite-activity) case, where the burst mechanism is parameterized directly by an arrival rate *κ* and a jump-size law *Y* for which we obtain:

##### Theorem 1

**(Stationary transform, moment extraction, and invertibility)** *In the compound-Poisson case with Π*(*dz*) = *κv*(*dz*) *and jump-size law Y*∼ *v, writing the jump term in the equivalent uncompensated compound-Poisson form described in Supplementary Material, Theorem 4, the stationary Laplace transform* 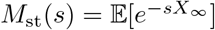 *is*

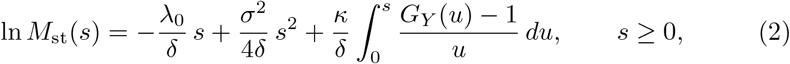

where *G*_*Y*_ (*s*) = 𝔼[e^−*sY*^ ]. *If μ_1_* = 𝔼[*Y*] *and μ_1_* = 𝔼[*Y*^*2*^] < ∞, *then*

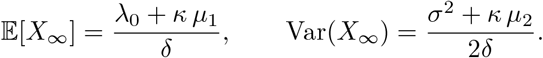

*Moreover*,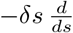ln 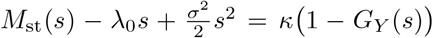, *so, once* (*λ*_0_, *δ, σ, κ*) *are fixed, the stationary transform uniquely determines the burst-size transform G*_*Y*_ .

The proof of Theorem 1 and the infinite-activity extension are given in Supplementary Material, Section A.4.

#### Efficient generation of traces

Expressing the surrogate models within the unified SDE framework directly allows one to vectorize generation of traces: the SDE is discretized into update steps of a fixed time step *Δt >* 0, iteratively applying deterministic, Gaussian, and jump-noise terms to the current state. Instead of updating a single trajectory, states can be updated as vectors of batch sizes, leading to the generation of a large number of trajectories simultaneously. Towards that goal we implemented the `bcrnnoise` framework so that deterministic and noise components support vector formats. For the update of the deterministic component we use basic and efficient first-order Euler integration; higher-order methods or intermediate substeps, while possible, would add significant computational cost and unbalance the treatment of noise and deterministic components with different step sizes. All subsequently discussed models make use of vectorized updates (see Methods for details).

We next discuss surrogate models for the one-state and the two-state model with respect to parameterization, computational cost, and dimensionality of the model’s state space; see Supplementary Material, Table 1 for an overview.

**Table 1:** Overview of the stochastic gene-expression models used in this work.

| Model | Symbol | States | Cost per $dt$ | Drift / noise |
| --- | --- | --- | --- | --- |
| *One-state models* |  |  |  |  |
| Exact birth-death (ref.) | $\mathcal{M}_{\text{CME}}$ | 1 | Gillespie (variable) | rates $\alpha n_p V$, $\delta x V$ |
| Tau-leaping | $\mathcal{M}_{\text{TL}}$ | 1 | 1 Pois + 1 Bin | thinned Poisson over $dt$ |
| CLE | $\mathcal{M}_{\text{CLE}}$ | 1 | 2 $\mathcal{N}$ | $(\alpha n_p - \delta x) dt + \sqrt{(\alpha n_p + \delta x)/V} dW$ |
| CLE (optimised) | $\mathcal{M}_{\text{CLE}^*}$ | 1 | 1 $\mathcal{N}$ | same SDE, single draw |
| Geometric bursts | $\mathcal{M}_{\text{Geo}}$ | 1 | 1 Pois + $K$ Geo | $-\delta x dt + \text{CPP}(\kappa, \text{Geom}(p))$ |
| Bin+Pois | $\mathcal{M}_{\text{BP}}$ | 1 | 1 Bin + 1 Pois | $M/M/\infty$ kernel; exact for any $\Delta t$ |
| *Two-state models* |  |  |  |  |
| Exact telegraph (ref.) | $\mathcal{M}_{\text{CME},2}$ | 3 | Gillespie (variable) | 4 reactions; $S_t \in \{0, 1\}$ |
| Tau-leaping | $\mathcal{M}_{\text{TL},2}$ | 3 | 4 Pois | 4 independent Poisson batches |
| CLE | $\mathcal{M}_{\text{CLE},2}$ | 3 | 4 $\mathcal{N}$ | per-reaction diffusion; $B_i^2(x) = f_i/V$ |
| CLE (optimised) | $\mathcal{M}_{\text{CLE}^*,2}$ | 3 | 2 $\mathcal{N}$ | mRNA block + switching block |
| Geometric bursts | $\mathcal{M}_{\text{Geo}}$ | 1 | 1 Pois + $K$ Geo | $\kappa, p$ matched to telegraph moments |
| Geo + analytical promoter | $\mathcal{M}_{\text{Geo},A}$ | 1 | 1 Pois + $K$ Geo | $N_{\text{on}}$ replaced by mean $\pi_{\text{on}} n_p (1 - e^{-(k_{\text{on}} + k_{\text{off}})t})$ |
| Bin+Pois | $\mathcal{M}_{\text{BP},2}$ | 3 | 3 Bin + 1 Pois | Kendall kernel for promoter; $M/M/\infty$ for mRNA |

### 3.2 Surrogates for one-state model

The following surrogate models approximate ℳ_CME_ using a single tracked variable (mRNA concentration) and fixed-size time steps. We start with the commonly used surrogates Tau-leaping and CLE.

#### ℳ_TL_: *Tau-leaping*

The model approximates ℳ_CME_ by combining reaction counts over a step *Δt*: mRNA births *N*_*b*_∼ Pois(*αn*_*p*_*V Δt*) and deaths *N*_*d*_∼Pois(*δxV Δt*) are chosen independently, and update *Δx* = (*N*_*b*_− *N*_*d*_)*/V* applied. Equivalently, one may draw the total number of events *N*∼ Pois((*αn*_*p*_ + *δx*)*V Δt*) and assign births by binomial thinning. The model recovers the SSA as *Δt*→ 0. *State:* 1 variable. *Cost per step:* 1 Poisson draw + 1 Poisson/Binomial draw.

#### ℳ_CLE_: *Chemical Langevin Equation*

Replaces the discrete reaction counts with their Gaussian fluctuation limit by applying the state update 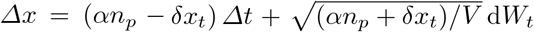. *State:* 1 variable. *Cost per step:* 2 Gaussian draws (one per reaction channel).

#### ℳ_CLE*_ : *Optimised CLE*

This is the first improved model. We collapse the two per-reaction Gaussian channels of ℳ_CLE_ into a single draw, exploiting the fact that birth and death noise are independent channels with their sum again being Gaussian with net diffusion coefficient 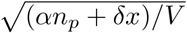. It is statistically identical to ℳ_CLE_, but improves runtime. *State:* 1 variable. *Cost per step:* 1 Gaussian draw.

#### ℳ_Geo_: *Geometric bursts*

We also tested a non-Gaussian model with heavier tails. While this model was primarily designed to capture bursts in the two-state model, we added it here for comparison. The model is a compound-Poisson surrogate with geometrically distributed burst sizes *Y* ∼Geom(*p*) arriving at rate *κ*. To improve computing time, we abstract fast mRNA decay into deterministic linear decay *f* (*x*) = −*δx*. The parameters *κ* and *p* are matched to the first two stationary moments of the reference model. *State:* 1 variable. *Cost per step:* 1 Poisson draw (burst count *K*) + *K* geometric draws; The model is expected to be fast if frequently *K* = 0 (i.e., bursts are rare). Since *K* ∼ Poisson(*κV Δt*), so 𝔼[*K*] = *κV Δt*, the geometric draws are rare whenever *κV Δt* ≪ 1.

#### ℳ_BP_: *Binomial–Poisson propagator*

Jahnke and Huisinga [22] gave a closed-form solution of the CME for monomolecular networks. Writing the one-step kernel in copy-number form with *X*_*t*_ = *V x*_*t*_, each of the *X*_*t*_ molecules present at time *t* survives to *t* + *Δt* with probability *e*^−*δΔt*^, giving Bin(*X*_*t*_, *e*^−*δΔt*^) survivors [25]. New births form a Poisson process of rate *αn*_*p*_*V* ; thinning by survival to *t* + *Δt* yields Pois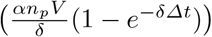. Hence, conditional on *X*_*t*_,

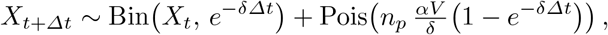

which is exact for any *Δt*. Unlike ℳ_TL_, which approximates production as Pois(*αn*_*p*_*V Δt*), ignoring concurrent degradation of newly produced molecules, and therefore has a one-step mean that grows without bound as *Δt*, ℳ_BP_ remains exact at any step size: as *Δt, e*^−*δΔt*^ → 0 so all current molecules degrade, and the Poisson mean converges to *n*_*p*_*αV/δ*, giving *X*_*t*+*Δt*_ ∼ Pois(*n*_*p*_*αV/δ*), the correct stationary distribution of the birth-death process, regardless of *X*_*t*_. *State:* 1 variable. *Cost per step:* 1 Binomial draw + 1 Poisson draw.

### 3.3 Surrogates for two-state model

We next discuss surrogates that approximate ℳ_CME,2_, again starting with the classical Tau-leaping and CLE models. The discussed models differ in how many variables they track and whether the promoter dynamics are simulated stochastically or collapsed analytically.

#### ℳ_TL,2_: *Tau-leaping*

The model extends ℳ_TL_ to the telegraph model by drawing four independent Poisson counts per step: transcription (*αN*_on_*V Δt*), degradation (*δxV Δt*), promoter off (*k*_off_ *N*_on_*V Δt*), and promoter on (*k*_on_*N*_off_ *V Δt*). *State:* 3 variables (mRNA, *N*_on_, *N*_off_). *Cost per step:* 4 Poisson draws.

#### ℳ_CLE,2_*:Chemical Langevin Equation*

Extends ℳ_CLE_ to the telegraph model by applying the Gaussian fluctuation limit to each of the four reaction channels (transcription, degradation, promoter off, promoter on) independently, with perreaction diffusion coefficients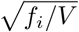. *State:* 3 variables (mRNA, *N*_on_, *N*_off_). *Cost per step:* 4 Gaussian draws.

#### ℳ_CLE*\**,2_: *Optimized CLE*

As in the one-state model, we collapses reaction noise channels of ℳ_CLE,2_: one for the mRNA birth–death block and one for the promoter switching block (applied with opposite sign to *N*_on_ and *N*_off_). The model is statistically equivalent to ℳ_CLE,2_. *State:* 3 variables (mRNA, *N*_on_, *N*_off_). *Cost per step:* 2 Gaussian draws.

#### ℳ_Geo_: *Geometric bursts*

As in the one-state benchmark, we consider a compound-Poisson surrogate with geometrically distributed burst sizes *Y* ∼Geom(*p*) and deterministic mRNA degradation *f* (*x*) = − *δx*. In the two-state setting, the burst parameters are matched to the stationary mean and variance of the telegraph model, but promoter fluctuations are not tracked explicitly. This provides a one-dimensional heavy-tailed surrogate that reproduces the stationary moments of ℳ_CME,2_ while collapsing all promoter dynamics into an effective burst process.*State:* 1 variable (mRNA). *Cost per step:* 1 Poisson draw + *K* geometric draws.

#### ℳ_Geo,A_: *Geometric bursts with analytical promoter*

This model also uses geometric burst sizes and deterministic mRNA decay, but improves on ℳ_Geo_ by retaining the transient activation of the promoter pool. We reduce the two-state system to a single mRNA variable by replacing the stochastic promoter occupancy *N*_on_(*t*) with its analytical conditional mean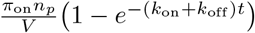,which captures the transient warm-up of the promoter pool when starting with all promoters in the off state. The geometric burst rate is then modulated proportionally to this deterministic envelope, while the burst-size parameter remains the stationary moment-matched one. *State:* 1 variable (mRNA). *Cost per step:* 1 Poisson draw + *K* geometric draws.

#### ℳ_BP,2_: *Binomial–Poisson propagator*

We also propose a new extension of ℳ_BP_ to the two-state telegraph model. Writing the one-step kernel in copy-number form with *X*_*t*_ = *V x*_*t*_ and *G*_*t*_ = *V N*_on,*t*_, we treat the *n*_*p*_ promoter copies as independent two-state Markov chains [25]. For a single promoter *S*_*t*_ ∈ {0, 1},

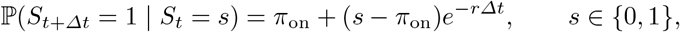

with *r* = *k*_on_ + *k*_off_ and *π*_on_ = *k*_on_*/r*. Hence, the exact endpoint on-probabilities over *Δt* are *p*_11_ = *π*_on_ +(1− *π*_on_)*e*^−*rΔt*^ and *p*_01_ = *π*_on_(1 −*e*^−*rΔt*^), and, conditional on *G*_*t*_,

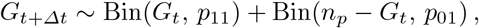

which is exact for any *Δt*. Freezing the promoter count at its left-endpoint value *G*_*t*_, the mRNA update is

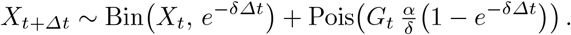

Unlike ℳ_BP_, whose constant birth rate makes the update exact without further conditioning, this mRNA step assumes the promoter count is constant over [*t, t* + *Δt*] and becomes exact as *Δt* → 0. Unlike binomial Tau-leaping [40,6], which uses the first-order birth approximation Pois(*αG*_*t*_ *Δt*), ℳ_BP,2_ accounts for degradation of molecules produced within the step: the new-survivor mean converges to *G*_*t*_*α/δ* as→∞*Δt*, the correct conditional stationary copy-number mean given the frozen promoter state. ℳ_BP,2_ therefore degrades more gracefully than other Tau-leaping variants at large *Δt. State:* 3 variables (mRNA, *N*_on_, *N*_off_). *Cost per step:* 3 Binomial draws + 1 Poisson draw.

We next derive analytic formulas for parameters for surrogate models where not already given by the initial parameters. The parameterized models were then benchmarked with respect to runtime and accuracy.

### 3.4 Model parameterization

Models are parameterized in one of two ways: they use the biological rates directly or they are parameterized by two-moment matching to the stationary mean and variance. We only state formulas for models that do not directly use biological parameters. Moment formulas are stated in concentration units.

For the implemented geometric-burst models, a time step *Δt* samples *K* ∼Poisson(*κV Δt*) burst events, each adding *Y/V* to the mRNA concentration, where *Y* ∼Geom(*p*) on ℕ_0_. Equivalently, in the notation of Theorem 1, the concentration jump law is *Y/V* and the arrival rate is *κV*. Hence, with *λ*_0_ = *σ* = 0, it is 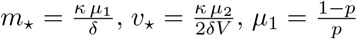,and 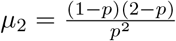.Solving for *p* and *κ* given a target (*m*_*_, *v*_*_) with 2*V v*_*_ *> m*_*_ gives

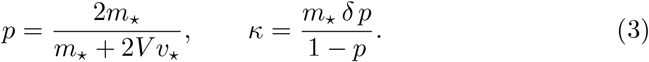

These formulas are instantiated with the one-state target moments below and, independently, with the two-state target moments.

#### Surrogates for one-state model

The stationary distribution of ℳ_CME_ with *n*_*p*_ constitutive copies is Poisson (Theorem 2), with concentration moments

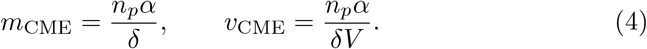

ℳ_CLE*_ : diffusion coefficient *B*^2^(*x*) = (*αn*_*p*_ + *δx*)*/V* .

ℳ_Geo_: Substituting Eq. (4) into Eq. (3) yields 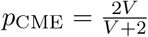and 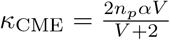At *V* = 1 fL: *p* = 2*/*3, *κ* = 2*n*_*p*_*α/*3. The feasibility condition *v*_CME_ *> m*_CME_*/*2 requires *V <* 2 fL.

#### Surrogates for two-state model

The stationary concentration moments of ℳ_CME,2_ with *n*_*p*_ promoter copies are (Theorem 3), with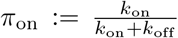 has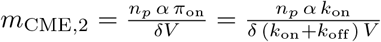and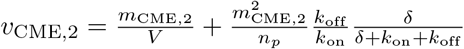.

ℳ_CLE*\**,2_: two block-wise diffusion coefficients from CME propensities, 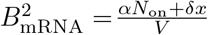 and 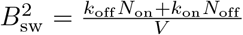

ℳ_Geo_: Substituting (*m*_*_, *v*_*_) = (*m*_CME,2_, *v*_CME,2_) into Eq. (3) yields 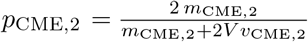 and 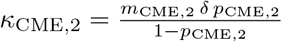

ℳ_Geo,A_: uses the same burst-size parameter *p*_CME,2_ and, because all promoters start in the off state, the analytical mean active-promoter concentration is 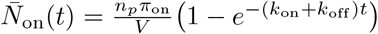.Its time-dependent burst propensity is there-fore 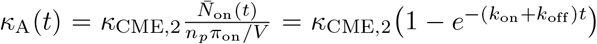 .The burst count over one step is sampled as *K* ∼ Poisson(*κ*_A_(*t*)*V Δt*).

## 4 Results

The parameterized models were then benchmarked with respect to runtime and accuracy after implementation within the `bcrnnoise` library.

### Runtime

We benchmarked runtime by simulating 100 trajectories of length 120 min for each model (30 repeats), varying the plasmid copy number *n*_*p*_ and, for surrogate models, the time step *Δt*, using literature-derived parameter sets (see Methods). For SDE-based surrogates, the 100 trajectories were generated in a vectorized batch. For the constitutive one-state model (Fig. 2a), all SDE-based surrogates have simulation times one to two orders of magnitude faster thanℳ _CME_. Unlike exact ℳ_CME_, their maximum runtime does not increase from *n*_*p*_ = 1 to 20 and is mainly determined by *Δt*. For the telegraph two-state model (Fig. 2b), similar observations hold. ℳ_BP_ and ℳ_BP,2_ are seen to perform similarly to ℳ_TL_ and ℳ_TL,2_. Geometric surrogates perform particularly well for the two-state model. Interestingly, ℳ_CLE*_ and ℳ_CLE *_,2 do not significantly improve performance of ℳ_CLE_ and ℳ_CLE,2_, which is attributed to the fact that random draws do not happen in the update loop but are generated in vectors upfront. Representative mRNA concentration trajectories for surrogate models of the constitutive one-state (Fig. 2c) and telegraph two-state (Fig. 2d) models are shown for the plasmid numbers *n*_*p*_ ∈ {1, 20} and the time-steps *Δt* = 1 min.

**Fig. 2:**
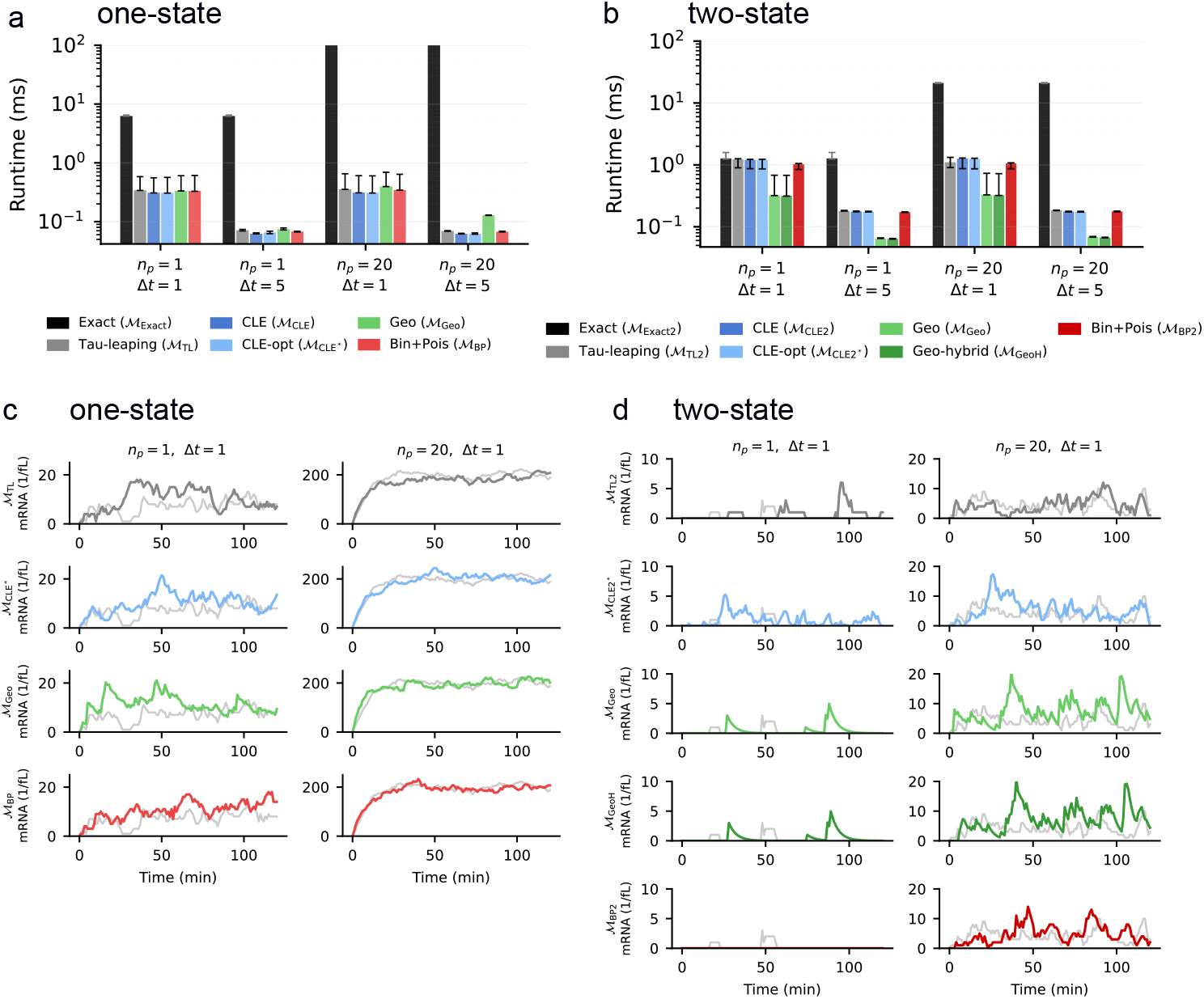
Runtime and trajectories for surrogate models. (**a**) Constitutive one-state model families benchmarked over 100 generated traces at *n*_*p*_ *∈{*1, 20} and *Δt∈{* 1 min, 5 min}. Bars show median runtime per trajectory across 30 repeated benchmark batches; error bars indicate the 16th–84th percentiles. (**b**) Telegraph two-state model families under the same conditions. (**c&d**) Example trajectories for surrogate models with reference trajectories from exact models shown (gray).

### Accuracy

Since surrogates achieved considerable performance boosts, we next tested their accuracy. We chose the same parameter settings as for the runtime benchmarks, generating 10,000 traces for each model to obtain precise noise profiles. Since ℳ_CLE_ and ℳ_CLE,2_ are identically distributed to ℳ_CLE*_ and ℳCLE_*_,2, we only show the optimized versions. The noise profiles were first tested in their accuracy to match marginal distributions of mRNA over the 120 min duration using Wasserstein-1 distance to the reference models at *Δt* time increments (Fig. 3a&b, see Methods for details). Early and late time distributions are also shown in the Supplementary Material (Section A.8).

**Fig. 3:**
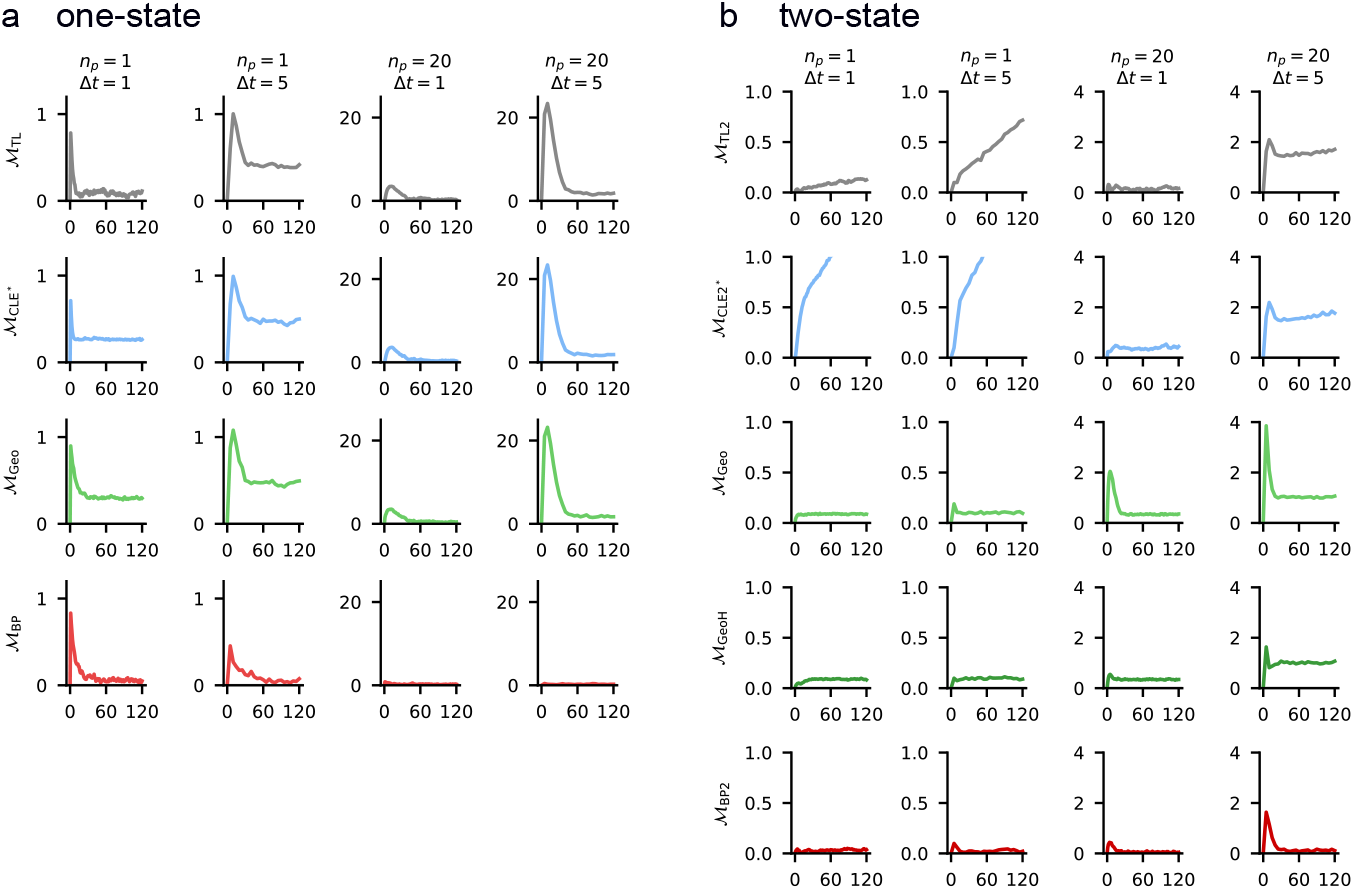
Wasserstein-1 distance over time for surrogate models. *W*_1_(*t*) between the marginal mRNA distribution of each surrogate and the exact reference (ℳ_CME_ for the constitutive one-state (**a**); ℳ_CME,2_ for the telegraph two-state (**b**)), estimated from 10,000 trajectories at each recorded time point. Rows: surrogate models; columns: plasmid copy number *n*_*p*_ *∈ {*1, 20*}* and time step *Δt ∈ {*1 min, 5 min*}*.

For the one-state model (Fig. 3a), at *n*_*p*_ = 1 and fast updates with *Δt* = 1 min, surrogates are seen to perform well, with ℳ_TL_ and ℳ_BP_ being the best. Increasing the step size to *Δt* = 5 min, however, considerably degrades accuracy except for ℳ_BP_. Similar behavior is observed for higher plasmid number *n*_*p*_ = 20, with the degradation for the early (transient) phase being large, except for ℳ_BP_. For the two-state model (Fig. 3b), ℳ_CLE*_ ,2 is seen to not capture the tailed distribution shapes with Gaussian noise and degrades even more over time. Geometric surrogates perform particularly well along with ℳ_BP,2_. Interestingly, increasing the step-size to *Δt* = 5 min does not lead to a large degradation of accuracy for geometric models; in contrast to ℳ_TL,2_ where the assumption of small propensity changes during *Δt* is not accurate anymore. Again, ℳ_BP,2_ shows excellent performance. For *n*_*p*_ = 20 plasmids, the transient phase, after switching on the promoters, is again the most challenging to approximate. The problem is pronounced for ℳ_Geo_, whose parameters are fitted to match the steady-state. This was alleviated by our model ℳ_Geo,A_ that accounts for the switching-on phase, considerably improving the accuracy.

We next tested the models’ capabilities to capture noise dependencies over time: noise at time *t* is not expected to be independent of noise at time *t*^′^ in a model with bursts. We thus computed normalized auto-correlation functions (ACFs) for all surrogate models and compared these to the exact models (see Methods for details). For the constitutive one-state model (Fig. 4a), all surrogates reproduce the reference ACF closely across both copy numbers for the small time-step of 1 min: the exponential decorrelation is captured by all models. For the larger time-step of 5 min, all models achieve acceptable correlation quality for *n*_*p*_ = 1, with the winner being the ℳ_BP_ model. Increasing plasmid counts to *n*_*p*_ = 20 visibly leads to strong underestimation of correlation except for the ℳ_BP_ model, which fits exactly the non-trivial correlation.

**Fig. 4:**
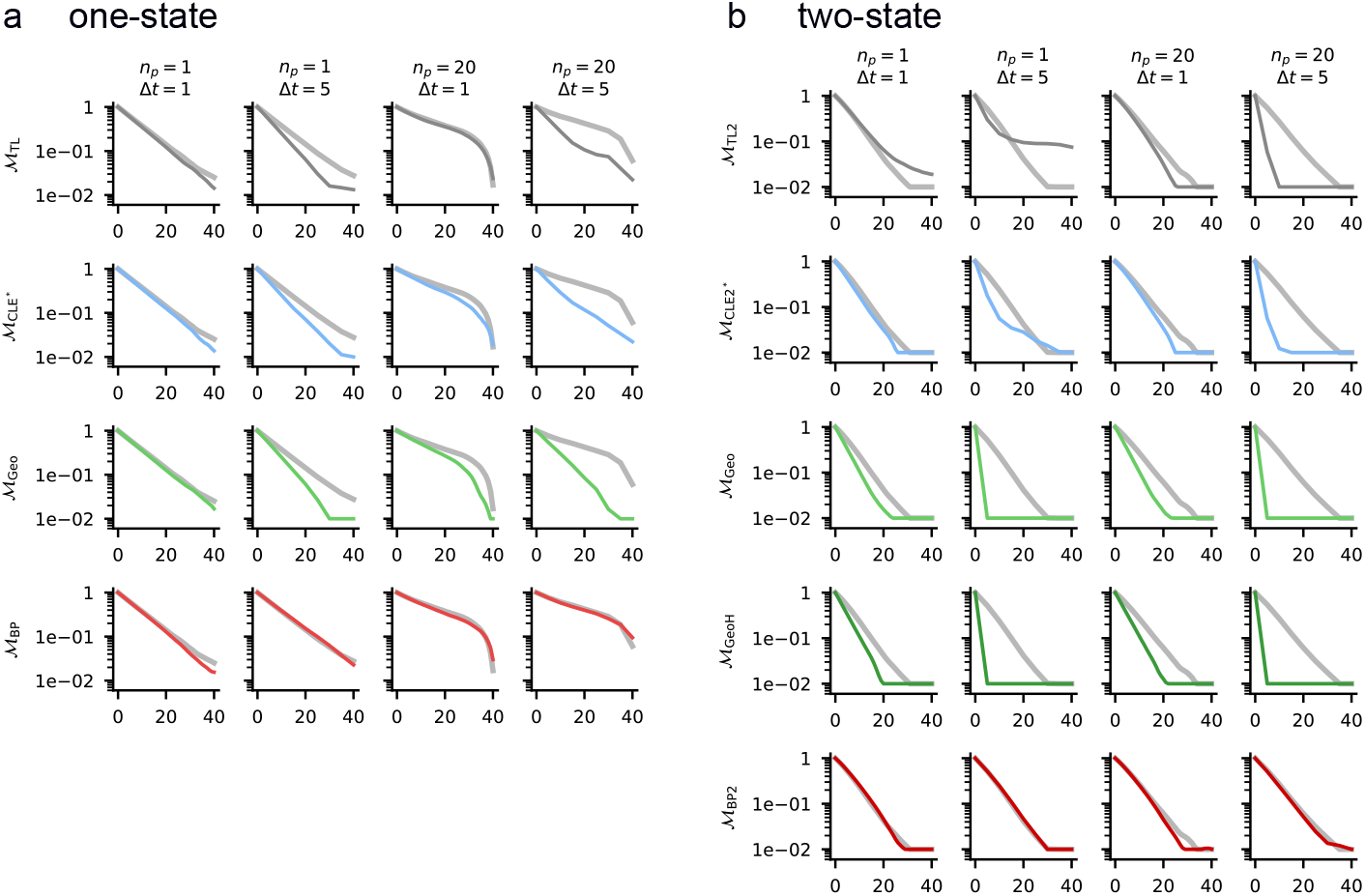
Autocorrelation functions of surrogate models. Empirical normalized ACF *ρ*(*τ*) of mRNA concentration for the constitutive one-state (**a**) and telegraph two-state (**b**) benchmarks, averaged over 10,000 trajectories. Rows: surrogate models; columns: plasmid copy number *n*_*p*_ 1, 20 and time step *Δt* 1 min, 5 min. The reference ACF (ℳ_CME_ or ℳ_CME,2_) is shown in gray in every panel; the surrogate ACF (log scale) is overlaid in the model color.

**Fig. 5:**
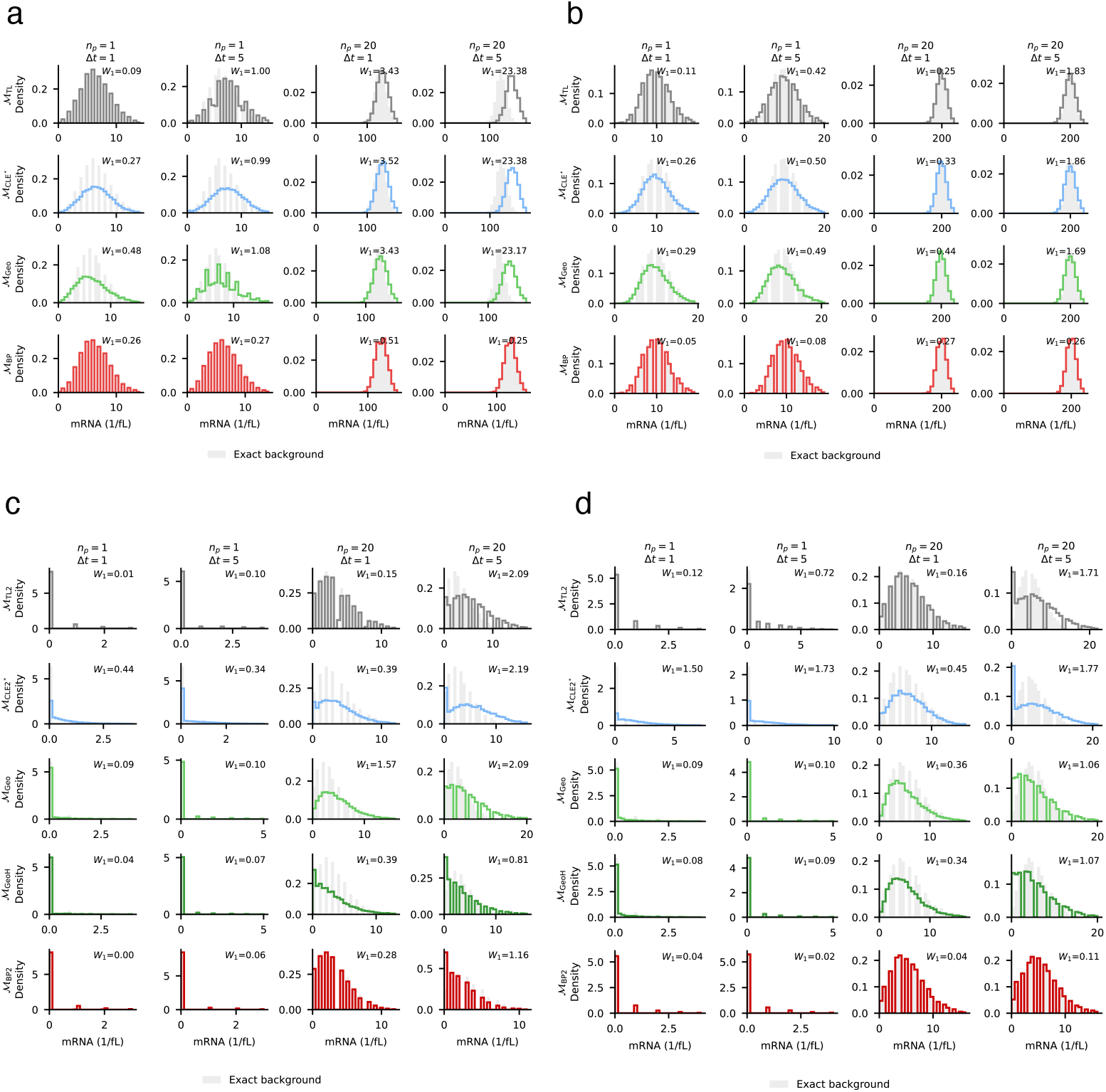
Marginal mRNA distributions at early and late times. Exact-reference and surrogate-model histogram comparisons for the constitutive one-state model at 10 min (**a**) and 120 min (**b**), and for the telegraph two-state model at 10 min (**c**) and 120 min (**d**). In each panel, rows correspond to surrogate models and columns to (*n*^*p*^, *Δt*) *∈* {(1, 1 min), (1, 5 min), (20, 1 min), (20, 5 min)}. The exact marginal distribution is shown as a gray background and the surrogate as a colored histogram outline.

For the telegraph two-state model (Fig. 4b), the short update-time of 1 min leads to favorable correlation performance for all models, with the geometric models moderately underestimation correlation. When increasing the update step to 5 min, large underestimations of the surrogate models become visible except for ourℳ_BP,2_ model. Interestingly, both one-dimensional geometric models struggle similarly to capture the slower correlation decline due to bursts within one dimension: ℳ_Geo,A_ does not improve over ℳ_Geo_ here, despite its superior prediction for marginals.

## 5 Methods

### Software implementation

All simulations were performed in our Python library `bcrnnoise` [10] available at https://github.com/BioDisCo/bcrnnoise. The library provides a unified interface for biochemical reaction network simulation under interchangeable noise specifications. A self-contained code listing illustrating this for a constitutive transcription network is given in Section A.6. All physical quantities (rates, concentrations, volumes, and time) are represented as `pint` [19] `Quantity` objects, providing automatic unit consistency checks and conversion throughout the library. We used the unit-jit library (https://github.com/BioDisCo/unit-jit) to just-in-time compile `pint` code to floats except at function boundaries. All random-number generation used NumPy. SDE-based models exploit a vectorized Euler–Maruyama kernel: all *n* trajectories advance simultaneously in a single loop iteration. For the compound-Poisson burst models (ℳ_Geo_, ℳ_Geo,A_), burst counts are drawn jointly across trajectories and the resulting geometric burst sizes are aggregated without pertrajectory loops using `np.add.reduceat`.

### Exact model parameters

*One-state [4]* Parameters are drawn from genome-wide median mRNA kinetics in *E. coli* (MG1655) [4]: per-plasmid transcription rate *α* = 1.0 fL^−1^ min^−1^, degradation rate *δ* = 0.1 min^−1^ (mean mRNA life-time 10 min). Cell volume was chosen as *V* = 1 fL. *Two-state [16]:* Parameters are from single-molecule measurements of the *λ* PRM promoter in *E. coli* [16]: *k*_on_ = 0.027 min^−1^ (mean OFF duration of 37 min), *k*_off_ = 0.17 min^−1^ (mean ON duration of 6 min), per-plasmid transcription rate *α* = 0.4 min^−1^, *δ* = 0.2 min^−1^ (mean mRNA lifetime 5 min), and *V* = 1 fL.

### Runtime metrics

Wall-clock times were measured with `time.perf_counter` over 30 batches of 100 trajectories of length *T* = 120 min on a single CPU core (MacBook Pro, Apple M2, 24 GB RAM). Exact reference models (ℳ_CME_, ℳ_CME,2_) used Gillespie simulation with variable step size.

### Accuracy metrics

Distributional accuracy was quantified by the first Wasserstein distance *W*_1_ using `scipy.stats.wasserstein_distance`. The empirical autocorrelation function (ACF) was computed from the same trajectories used for the distributional validation. To avoid transient effects, the first 40 min of each trajectory were discarded. The remaining stationary portion spans *t*∈ [40 min, 120 min]. For each model the unnormalized autocorrelation at integer lag *k* was estimated as 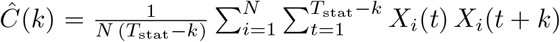,where *N* = 10,000 is the number of trajectories and *T*_stat_ is the number of retained time points. The ACF was then normalized to 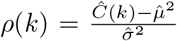, with 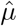 and 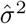the mean and variance over all trajectories and retained time points, so that *ρ*(0) = 1. Lags were evaluated up to 40 min.

## 6 Conclusion

We introduced a unified jump-diffusion SDE framework for mRNA trace generation in the constitutive and telegraph transcription models. The framework places classical approximations (Tau-leaping, the Chemical Langevin Equation) and compound-Poisson burst surrogates within a single drift–noise specification, and includes new surrogates based on geometric noise and Binomial–Poisson propagators. Implemented in the open-source `bcrnnoise` library, it enables vectorized batch simulation and comparison against exact Gillespie references.

Benchmarking shows that noise-model choice matters greatly for accuracy and efficiency. Gaussian surrogates are fast but lose accuracy in burst-dominated regimes, particularly for the telegraph model and during transient activation. Geometric burst surrogates reproduce marginal distributions substantially better, and an analytical warm-up of promoter activation further improves transient behavior, however perform poorly to capture correlation.

Most notably, Binomial–Poisson propagators combine high accuracy with strong computational performance, remaining accurate where Tau-leaping degrades at coarse time steps. The resulting two-orders-of-magnitude speed-up at near-identical accuracy provides a practical basis for selecting surrogates by observable and biological regime.

## Acknowledgments

The research of Arman Ferdowsi was funded in part by CHIST-ERA-22-SPiDDS-07 (TROCI Project), by the Austrian Science Fund (FWF) under grant 10.55776/I6647, and by the Austrian Science Fund (FWF) project DMAC (grant 10.55776/P32431). The research of Matthias Függer and Thomas Nowak was supported by the French National Research Agency (ANR) projects DREAMY (ANR-21-CE48-0003) and COSTXPRESS (ANR-23-CE45-0013), as well as the SAIF project, funded by the “France 2030” government investment plan managed by ANR, under the reference ANR-23-PEIA-0006.

## A Supplementary Material

### A.1 Stationary distribution for the constitutive one-state model

The constitutive one-state model steady-state follows a Poisson distribution [39,32]. We give a self-contained proof for completeness.

#### Proposition 2

*Consider the continuous-time birth-death Markov chain on ℕ*_0_ *with constant birth rate a* = *αn*_p_*V >* 0 *and death rate δn from state n, where δ >* 0. *Its unique stationary distribution is*

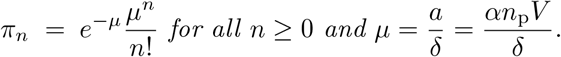

*That is, the stationary mRNA copy number is* Poisson(*αn*_p_*V/δ*).

*Proof*. Set *µ* = *a/δ* and define

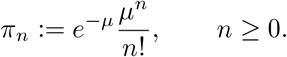

First,

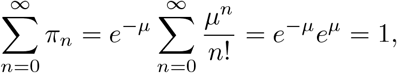

so (*π*_*n*_)_*n*≥0_ is a probability distribution.

It remains to check stationarity. For every *n* ≥ 0,

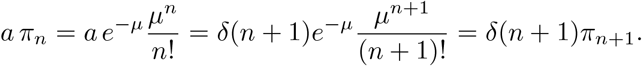

Thus the local balance relation *a π*_*n*_ = *δ*(*n* + 1)*π*_*n*+1_ holds for all *n* ≥0.

Let *Q* be the generator of the chain. For *n* = 0, (*πQ*)_0_ = −*aπ*_0_ + *δπ*_1_ = 0. *n* ≥1, (*πQ*)_*n*_ = *aπ*_*n*-1_ + *δ*(*n* + 1)*π*_*n*+1_ − (*a* + *δn*)*π*_*n*_ = *δnπ*_*n*_ + *a π*_*n*_ − (*a* + *δn*)*π*_*n*_ = 0, where we used the local balance relation twice: *aπ*_*n-*1_ = *δnπ*_*n*_ and *δ*(*n* + 1)*π*_*n*+1_ = *aπ*_*n*_. Hence *πQ* = 0, so *π* is stationary.

The chain is irreducible on *ℕ*_0_: from any state one can reach any other state by a finite sequence of births and deaths, each with positive rate. An irreducible countable-state continuous-time Markov chain has at most one stationary probability distribution. Therefore *π* is the unique stationary distribution.

### A.2 Exact moments for the telegraph two-state model

The stationary mean and variance of the telegraph model are given in [34,35]. We state a self-contained proof for completeness.

#### Proposition 3

*Consider n*_p_ *independent promoter copies. For each i* ∈ {1, …, *n*_p_}, *let* 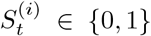 *switch at rates k*_on_ *(off*→*on) and k*_off_ *(on*→*off). When* 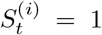 = 1, *promoter i produces mRNA at rate α, and each mRNA molecule degrades at rate δ. Let X*_*t*_ *be the total mRNA copy number and set*

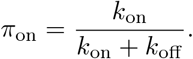

*Then the process admits a unique stationary distribution, and its stationary moments are*

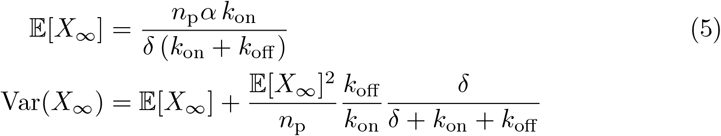

*Proof*. We first treat the single-promoter case *n*_p_ = 1. Then (*X*_*t*_, *S*_*t*_) ∈ ℕ_0_ × 0, 1 is a continuous-time Markov chain with generator

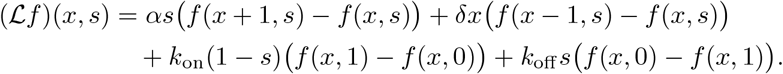

The chain is irreducible. Moreover, with *V* (*x, s*) = *x*,

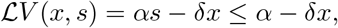

and with *V*_2_(*x, s*) = *x*^2^,

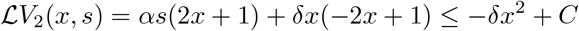

for some finite constant *C*. These drift bounds imply positive recurrence and finiteness of the stationary second moment. Hence a unique stationary distribution exists, and for the test functions used below we may write

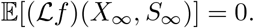

Taking *f* (*x, s*) = *s* gives

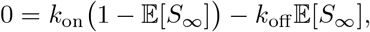

so

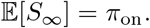

Taking *f* (*x, s*) = *s* gives

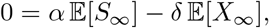

hence

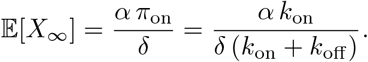

Now set *C* := 𝔼 [*X*_∞_*S*_∞_]. Taking *f* (*x, s*) = *xs* yields

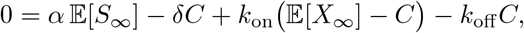

therefore

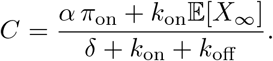

Finally, taking *f* (*x, s*) = *x*^2^ gives

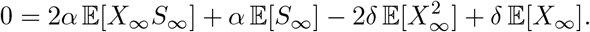

Using *δ* 𝔼[*X*_∞_] = *απ*_on_, this becomes

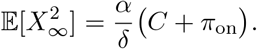

Substituting the expressions above and simplifying gives, for one promoter copy,

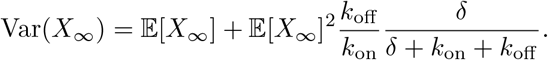

For general *n*_p_, label each transcript by its promoter of origin. Then the total mRNA count can be written as

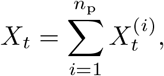

where the *n*_p_ processes 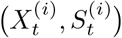 are independent copies of the single-promoter model. If *m*_1_ and *v*_1_ denote the stationary mean and variance for one promoter, then by independence

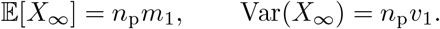

Using the one-promoter formulas 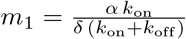, and 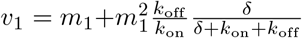 we obtain

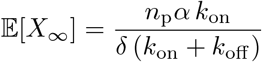

and

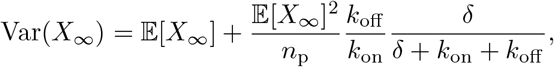

as claimed.

If *x*_*t*_ := *X*_*t*_*/V* denotes the mRNA concentration, then the corresponding stationary moments are

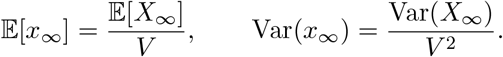

*One-step promoter kernel for ℳ*_BP,2_. For a single promoter *S*_*t*_ ∈ {0, 1}, let

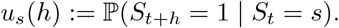

Then

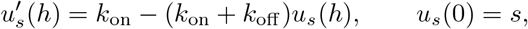

so with *r* = *k*_on_ + *k*_off_ and *τ*_on_ = *k*_on_/*r*,

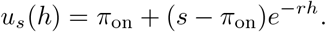

Hence

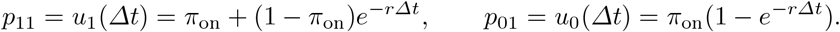

Conditional on *G*_*t*_ active promoter copies at time *t*, independence of the *n*_*p*_ promoters yields

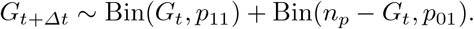

### A.3 Properties of unified jump-diffusion SDE

#### Theorem 4

**(Uniqueness and positivity)** *Let X*_*t*_ ∈ ℝ*d solve the Lévy–Itô SDE*

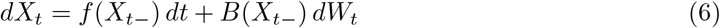

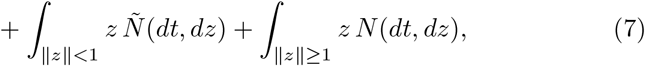

*where W*_*t*_ *is a d-dimensional Brownian motion and N* (*dt, dz*) *is a Poisson random measure on* ℝ_+_ × (ℝ*d*\{ 0}) *with intensity dt Π*(*dz*), *with Ñ* (*dt, dz*) = *N* (*dt, dz*) − *dt Π*(*dz*) *on* {∥*z* ∥ *<* 1}.

Uniqueness. *Assume that f and B are locally Lipschitz and satisfy a standard non-explosion condition (e*.*g. a linear-growth bound), and that the Lévy measure Π satisfies*

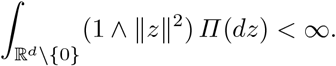

*Then there exists a unique strong càdlàg solution on* [0, ∞).

Positivity. *Suppose B* ≡ 0, *Π is supported on* 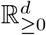, *and either the jump input is compound Poisson or, more generally*,

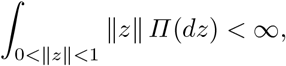

*so that the small-jump compensator can be absorbed into the drift and the equation rewritten as*

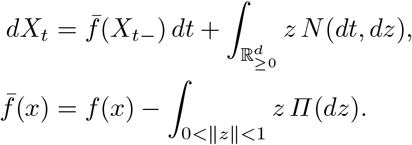

*If, for each i* ∈ {1, …, *d*}, *the boundary condition*

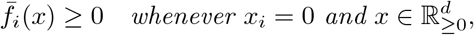

*holds, then the process remains in* 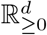 *for all t* ≥ 0 *almost surely for all non-negative initial data*.

*Proof. [sketch]* Existence and pathwise uniqueness are standard for Lévy–Itô SDEs under local Lipschitz assumptions together with a non-explosion condition; see, for example, [1, Thm. 6.2.3].

For positivity, the relevant object is the uncompensated nonnegative jump representation: once the jump term is written as 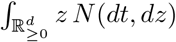, jumps can-not leave the nonnegative orthant, and the inward-pointing condition on 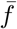 is the standard Nagumo-type viability condition [2]. By contrast, support of *Π* on 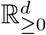 does *not* by itself imply positivity in the compensated representation 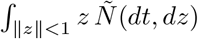, because the compensator contributes an additional finite-variation term.

One can further derive Kolmogorov forward equations for the density *p*(*x, t*) over species concentration vectors *x* and time *t*.

#### Theorem 5

**(Nonlocal Kolmogorov forward equation)**

*If p*(*x, t*) *is the density of Eq. (6), then*

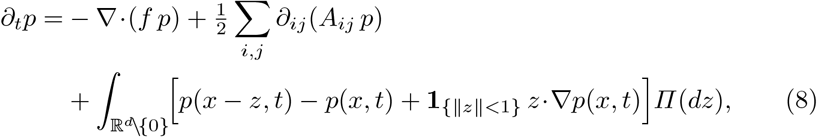

*where A*(*x*) = *B*(*x*)*B*(*x*)^⊤^.

*For the finite-activity compound-Poisson representation*

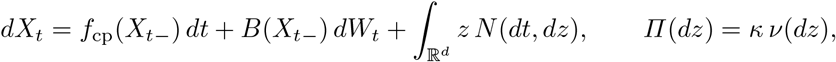

*equivalently* 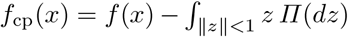*when starting from the compensated form, the forward equation reduces to*

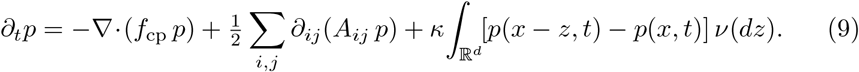

*Proof*. Let 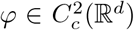 and write the generator of Eq. (6) on *φ* (Section A.5). Duality 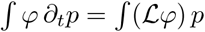 and integration by parts for the drift/diffusion terms yield the first two terms in Eq. (8); the jump term comes from the Lévy integral with compensation on{∥z∥<1}.For finite *Π*, rewrite the jump input in uncompensated compound-Poisson form after absorbing ∫_∥z∥<1_ *z Π*(*dz*) into the drift; a change of variables then gives Eq. (9).

### A.4 Properties of linear one-dimensional SDE

The transforms below are classical for linear SDEs driven by Lévy noise [1, Ch. 17]; for generalized Ornstein–Uhlenbeck stationarity see also [30].

#### Proof (Proof of Theorem 1 and infinite-activity extension)

For the finite-activity compound-Poisson form, multiply the forward equation Eq. (9) by *e*^−*sx*^ and integrate to obtain the PDE for *M* (*s, t*) = 𝔼[*e*^−*sX*^*t* ]:

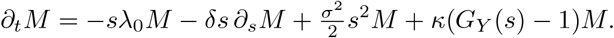

Solving the stationary first-order ODE in *s* yields Eq. (2). The moment formulas follow by differentiating ln *M*_st_ at *s* = 0, and the invertibility identity is obtained by rearranging the same ODE.

For general infinite-activity *Π*, apply the generator to *φ*(*x*) = *e*^*ikx*^, derive the stationary transport equation for *φ*(*k, t*) = 𝔼[*e*^*ikX*^*t* ], and integrate along characteristics to obtain the stationary characteristic function

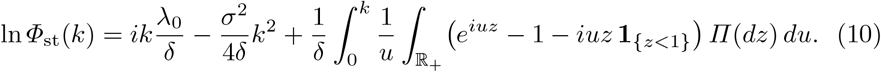

The OU/Lévy series representation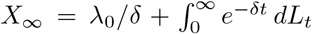is a standard equivalent formulation; see [1, Ch. 17].

The following result is a standard consequence of Karamata’s Tauberian theorem and tail-transfer for regularly varying laws in OU-type shot-noise processes [9, Eq. (4.18)].

##### Theorem 6

**(Tail transfer under heavy-tailed bursts)** *If either the driving subordinator has Laplace exponent ϕ*(*s*) ∼*c s*^*α*^ *as s*Φ 0 *with* 0 *< α <* 1, *or, in the finite-activity compound-Poisson case, the burst-size law satisfies* 𝕡{*Y > y*} ∼ *C y*^−*α*^, *then the stationary law satisfies*

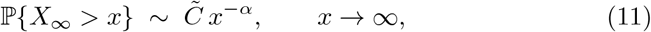

*Proof*. The stable-like infinite-activity case follows directly from the stationary Laplace exponent formula: if the driving subordinator has *ϕ*(*s*) ∼ *s*^*α*^ as *s* Φ0 with 0 *< α <* 1, then

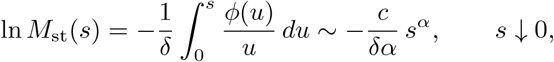

and Karamata’s Tauberian theorem yields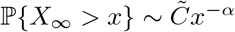.

The deterministic shift *λ*_0_*/δ* and, when *σ >* 0, the Gaussian OU component are light-tailed and therefore do not affect the regular-variation index. It is therefore enough to analyze the jump component. For the finite-activity compound-Poisson case with regularly varying burst-size tail 𝕡 {*Y > y}*∼ *{C y*^−*α*^*}*, it is cleaner to use the shot-noise / perpetuity representation

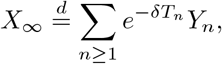

where (*T*_*n*_) are the jump times of a rate-*κ* Poisson process and the *Y*_*n*_ are i.i.d. copies of *Y*. Because regularly varying laws are subexponential, classical tail-transfer results for weighted sums and OU-type shot-noise processes imply that the stationary law has the same tail index *α*, up to a multiplicative constant depending on (*κ, δ*). This gives Eq. (11).

### A.5 Generator and the forward (Fokker–Planck) equation

For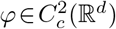the infinitesimal generator of Eq. (6) is

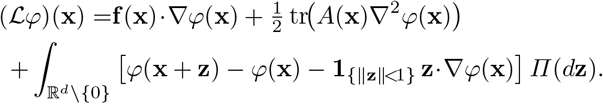

Duality against test functions yields Eq. (8). When *Π* is finite, one may rewrite the jump input in uncompensated compound-Poisson form after absorbing∫∥_z_ ∥_<1_**z** *Π*(*d***z**) into the drift. In that representation, Eq. (9) follows after a change of variables.

### A.6 Software and reproducibility: bcrnnoise usage example

To keep the main text concise while retaining reproducibility, we collect here a minimal schematic `bcrnnoise` example showing how the same transcription BCRN can be simulated either as an exact Markov jump process (SSA) or as an SDE by changing only the noise specification. For example, the BCRN comprising the species mRNA and the reactions∅→mRNA and mRNA→∅following mass-action kinetics with rate constants *α* (min^−1^ fL^−1^) for transcription and *δ* (min^−1^) for degradation/dilution, is specified in `bcrnnoise` via:

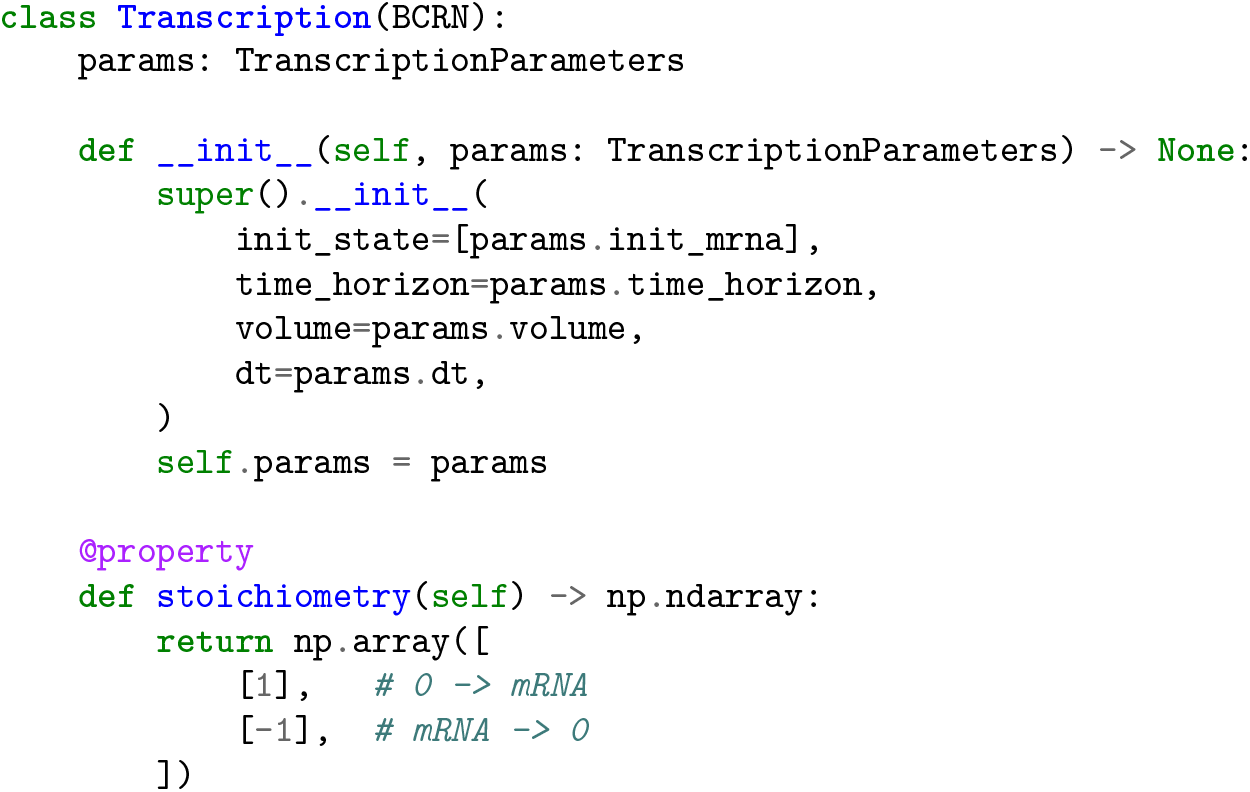
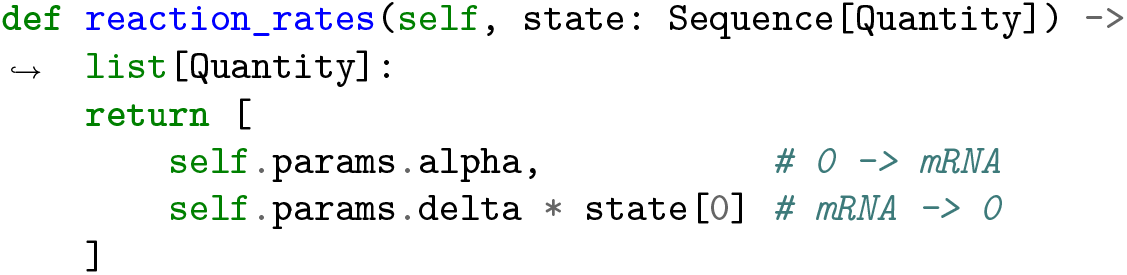

An exact SSA trajectory is then obtained via:

~~~
transcription = Transcription(params)
ts = transcription.simulate_markov_chain(seed=42)
~~~

To simulate the same system with Geometric noise for mRNA generation and deterministic degradation/dilution, one can change the above class to:

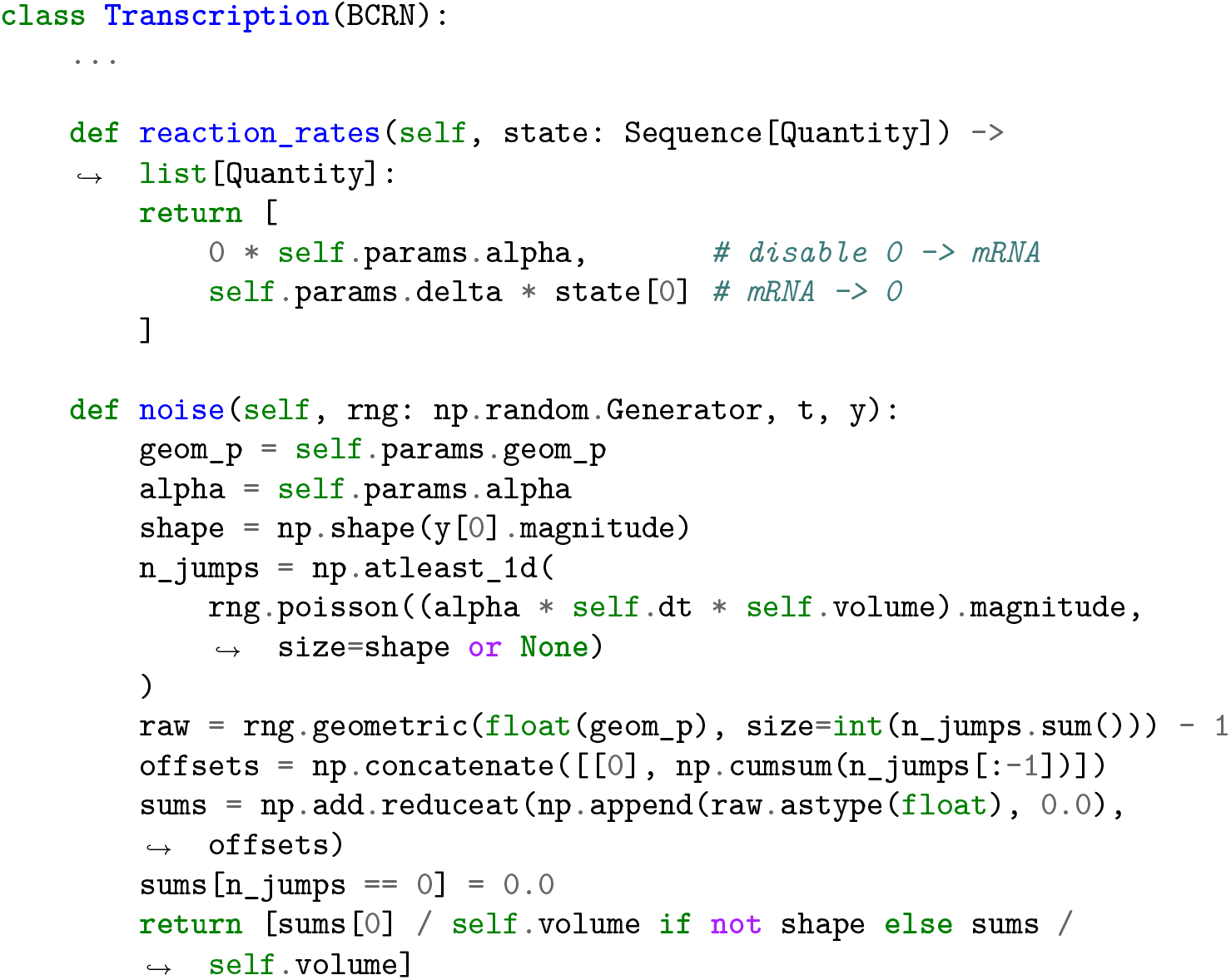

Simulation of the SDE is then performed via:

~~~
ts = transcription.simulate_sde(seed=42)
~~~

and vectorized batch simulation via:

~~~
batch = transcription.simulate_sde_batch(n=100, seed=42)
~~~

The library supports physical units via pint [19] and the same interface extends to other surrogate models used throughout Section 4.

### A.7 Model overview

Table 1 summarizes the state dimension, noise structure, and per-step stochastic cost of all models considered in this work. Here, *States* counts the number of tracked variables per trajectory, and *Cost per dt* counts random draws per step (Pois = Poisson, Geo = geometric,𝒩 = Gaussian); *K* ∼Pois(*κV dt*) denotes the realised burst count. All continuous-time models fit within the unified Lévy–Itô specification in Eq. (1).

### A.8 Supplementary Histogram Panel

